# Tradeoffs of increasing temperatures for the spread of antimicrobial resistance in river biofilms

**DOI:** 10.1101/2023.09.08.556853

**Authors:** Kenyum Bagra, David Kneis, Daniel Padfield, Edina Szekeres, Adela Teban-Man, Cristian Coman, Gargi Singh, Thomas U. Berendonk, Uli Klümper

**Affiliations:** Institute for Hydrobiology, Technische Universität Dresden, Dresden, Germany; Indian Institute of Technology, Roorkee, Uttarakhand, India; Environment and Sustainability Institute, University of Exeter, Penryn Campus, Penryn, UK; Institute of Biological Research Cluj, NIRDBS, ClujlZlNapoca, Romania

**Keywords:** Antimicrobial resistance, Invasion, River Biofilm, Climate Change, Temperature

## Abstract

River microbial communities regularly act as the first defense barrier against the spread of antimicrobial resistance genes (ARG) that enter environmental microbiomes through wastewaters. However, how the invasion dynamics of wastewater-born ARGs into river biofilm communities will shift due to increasing average and peak temperatures worldwide through climate change remains unknown. Here we aimed at elucidating the effects of increasing temperatures on both, the natural river biofilm resistome, as well as the river biofilms invadability by foreign, wastewater-born ARGs. To achieve this, natural biofilms were grown in a pristine German river and transferred to artificial laboratory recirculation flume systems at three different temperatures (20°C, 25°C, 30°C). Already after one week of acclimatization to the temperatures, significant increases in the abundance of most naturally occurring ARGs were detected in the biofilms exposed to the highest temperature. Thereafter, biofilms were exposed to a single pulse of wastewater and the invasion dynamics of wastewater-born ARGs were analyzed over a period of two weeks. While initially after one day ARGs were able to invade all biofilms successfully and in equal proportions, the foreign invading ARGs were lost at a far increased rate at 30°C over time. ARG levels dropped to the initial natural levels at 30°C after 14 days. Contrary at the lower temperatures ARGs remained far elevated and certain ARGs were able to establish themselves in the biofilms. Overall, we here demonstrate tradeoffs of increasing temperature between increases in naturally occurring and faster loss dynamics of invading ARGs.

## Introduction

The rise in antimicrobial resistance (AMR) on a global scale has been considered a silent pandemic (Mahoney et al., 2021). Based on the clinical reports available globally, 4.95 million people succumbed to bacterial infections with resistant pathogens in 2019 (Murray et al., 2020), with numbers expected to rise up to 10 million lives per year by 2050 (O’Neill, 2016). Mitigating the spread of AMR is difficult as there is no clear barrier between the environment, animal and human microbiomes, thus, it needs to be approached within a “One Health”-framework (Hernando-Amado et al., 2019). The increased risk of pathogens acquiring resistance to the limited range of clinically used antibiotics makes it imperative that we find ways to stop the spread of AMR within and between the three aforementioned microbiomes (Tacconelli et al., 2018).

Rivers and river biofilms could play a major role in the spread of AMR as rivers are not only one of the main sources of our daily water supply, but, in many parts of the world, they also serve as a conduit of waste and wastewater from households, hospitals, animal and aquatic farms (Berendonk et al., 2015; Hannah et al., 2022; Heß et al., 2018; Huijbers et al., 2019; Seiler and Berendonk, 2012). High loads of antibiotic-resistant bacteria (ARBs) and antibiotic-resistant genes (ARGs) are constantly transferred via fecal matter through wastewater streams to wastewater treatment plants (WWTPs) (Karkman et al., 2019). In wastewater, ARBs and ARGs are also exposed to chemical pollutants like pharmaceuticals, pesticides, and heavy metals, which induce selection pressure and promote horizontal gene transfer, thus making WWTPs a hotspot for the spread of AMR (Gupta et al., 2023; Li et al., 2015; Rizzo et al., 2013). Despite a general reduction in total bacterial numbers through WWTPs, the discharge of treated wastewater into rivers still introduces significant loads of ARGs and ARBs to the receiving waters (Ju et al., 2019; Zhang et al., 2021). Additionally, in many locations globally, wastewater is discharged without any prior treatment (Marathe et al., 2017; Narain, 2012; Williams et al., 2019) accounting for around 48% of the globally produced wastewater that enters environmental waters untreated (Jones et al., 2008). Even in countries like Japan, with one of the highest WWTP coverage in the world of 91.4%, untreated water from around 14 million people is still discharged into the environment (Baba et al., 2022). Moreover, extreme weather events like floods, heavy rainfall, monsoon and cyclones result in the overflow of untreated wastewater which is then released into the environment alongside stormwater (Ahmed et al., 2018; Garner et al., 2017; Hamilton et al., 2020; Huijbers et al., 2019).

In this context, river microbial communities regularly act as the first line of defense against the spread of ARB and ARG that, after enrichment in the anthroposphere, enter the environment through wastewaters (Bürgmann et al., 2018). Unsurprisingly, such discharges of treated and untreated wastewater are often directly linked to an increased level of AMR in the microbiomes of the receiving water bodies making it a crucial pathway for the dissemination of ARGs (Berendonk et al., 2015; Pruden et al., 2012). Consequently, understanding what drives the invasion success of ARB and ARG from wastewater into river microbiomes is essential to combat their establishment in river microbiomes.

When wastewater enters rivers, the stress caused by the release of chemical pollutants into the environment can directly impact the structure of microbial communities (Gupta et al., 2023; Wu et al., 2019). Disturbances of the microbial community structure can pave the way for the successful establishment of incoming resistant bacteria into river biofilm communities (Bagra et al., 2023). Exposure to stress is hence a defining factor in shaping the natural invasion barriers of environmental communities. One of the main stress factors that environmental microbiomes worldwide are experiencing currently is the increase in average and peak temperatures through climate change. According to the Intergovernmental Panel on Climate Change, the global temperature increased by 1°C since the pre-industrial era and it is expected that global temperature will rise by more than 1.5°C from 2030-2052 (IPCC, 2018).The rise in air temperature is likely to be accompanied by an increase in surface water temperatures as both are directly linked (Mantua et al., 2010; van Vliet et al., 2011). Many studies have reported shifts in microbial community composition and function in soils, river waters and sediments with such changes in temperature (Balser and Firestone, 2005; García-Armisen et al., 2014; Mai et al., 2018; Wang et al., 2023).

With regards to AMR in river microbiomes such a raise in temperature could result in two main outcomes: First, changes in community composition due to changing temperature could directly lead to the enrichment or loss of ARG host bacteria, depending on their adaptation to the warming conditions. Second, warming could also affect how river microbial communities function as a barrier to invading ARBs from wastewater. During such coalescence events the interplay between different temperature optima of the resident community and the resistant invaders could affect their competitive interactions and hence the outcome of the invasion events (Mansour et al., 2018).Furthermore, higher temperatures are regularly connected to higher metabolic and growth rates (Balser et al., 2006), which could intensify competitive interactions during the invasion events. Based on these predictions, we here determined if climate change does indeed affect AMR in river microbial communities by investigating temperature effects on both, the intrinsic river biofilm resistome as well as the outcome of invasion events of ARB and ARG from wastewater into these biofilms.

To achieve this, river biofilms were grown on glass slides which were immersed into a pristine sub-tributary of river Elbe in Germany for one month. The biofilms were then transferred to artificial laboratory recirculation river water flume systems. These flume systems were kept at three different temperatures of 20°C, 25°C and 30°C. Biofilms were allowed to acclimatize for one week at the respective temperature. Thereafter, biofilms were exposed to a single pulse of wastewater for 2 weeks. Throughout the experiment biofilms were destructively sampled and the biofilm resistome was characterized using high throughput qPCR, while the biofilm microbial community composition was analyzed using 16S rRNA gene-based amplicon sequencing. This allowed determining the time-resolved invasion dynamics of wastewater-born ARB and ARGs into the river biofilms as a function of temperature. In addition, control group flumes without wastewater addition were run to determine the effect of increasing temperatures on microbiome and resistome composition.

## Material & Methods

### Biofilms and river water

Natural river biofilms were obtained from river Hirschbach (50°54 ‘17.6“N, 13°45’ 06.3“E) which is a part of the Lockwitzbach tributary in Saxony, Germany. The chosen location had no WWTP or agricultural fields located upstream and was hence of low anthropogenic impact. Biofilms were grown on rectangular glass slides of 76 x 26 mm size (DWK Life Science, Wertheim, Germany) that were mounted on Artificial Exposure Units (AEUs). AEUs were constructed from Plexiglas (Fig. SI1). 12 AEUs each containing 9 glass slides were screwed onto a concrete plate and covered with a stainless-steel cover for protection and to provide shade to limit the growth of phototrophic organisms. A metal mesh at the front and the back allowed river water to stream through the AEUs. AEUs were immersed in the river for 48 days in March 2022 to allow biofilm growth. The AEUs containing the glass slides with grown biofilms were then collected and individually transferred to the laboratory in a 1L box containing river water at ambient temperature. Simultaneously, 100 L of the same river water was collected, immediately transported to the laboratory and filter sterilized using 0.2 µm pore size membrane filters (Whatman, Maidstone, UK) for use in the artificial flume systems.

### Laboratory flume system

Laboratory artificial flume systems (Bagra et al., 2023) with a volume of 40 x 19.1 x 10 cm (L x W x H) were constructed from polypropylene with an inlet and an outlet attached to each end (Fig. SI1). The inlet and outlet of the flume were connected to a 2L reservoir which was attached to a recirculation pump. Flumes were filled with a total volume of 6.348 L sterile river water constantly recirculated with the pump at 100 mL/min. The inlet was 1cm in diameter and at 0.5 cm height, a PVC pipe was inserted on the inner side of the system pointing towards the bottom of the flume to allow for an even distribution of the waterflow inside the flume system. The outlet was at the back, located at 7 cm height to maintain a constant maximum water level inside the flume. The artificial flumes were kept covered with polypropylene lids to avoid light entering the system. Three AEUs containing glass slides retrieved from the river were placed in the middle of each flume filled with sterile filtered river water from the site.

### Wastewater from the inlet of WWTP

Wastewater was collected from the influent of the WWTP located at Dresden-Kaditz, Germany (51.07 °N, 13.67 °E) on the 21st of March 2022, in a sterile 10L bottle. Suspended solids and sand particles were allowed to settle for 2 minutes and the resulting supernatant was used for flume experiments after any large debris still floating in the wastewater that could clog the flume system was removed manually with sterilized tweezers. For subsequent molecular analysis, 50 mL of wastewater was filtered in triplicate with 0.2 µm pore size membrane filter paper (Whatman, Maidstone, UK) and the filters kept at –20°C for downstream DNA extractions and analysis.

### Flume experiments

Two artificial flume systems containing AEUs with grown river biofilms were placed in each of the climate chambers set to 20°C, 25°C, and 30°C. The temperature was monitored throughout the experiment to remain constant. Before starting any treatments, the biofilms retrieved from the river site were allowed to acclimatize to the laboratory conditions for a week. After this period, one of the flumes from each chamber was inoculated with wastewater. A single pulse of 1.375 L (20% of flume volume) of influent was added while simultaneously removing excess water from the outlet. The other flume in each of the chambers were used as the control group where instead of the wastewater, 1.375 L of sterile river water was added. The flumes were then run for 14 days.

### Biofilm sampling

For each destructive sampling timepoint of biofilms, one individual glass slide with biofilm was randomly removed from each of the three AEUs of both the influent and control flume from each of the chambers, resulting in three biological replicates per timepoint and treatment. Each individual biofilm slide was carefully placed inside a 50 mL centrifuge tube with 1 mL of sterile detergent solution (0.9% NaCl solution with 0.05% Tween80® (Sigma-Aldrich, St. Louis, MO, USA)). The biofilms were scraped from the glass slide into the tube using a cell scraper (TPP, Trasadingen, Switzerland). The scraper and the glass slide were washed with the detergent solution to allow complete detachment of the biofilm from the scraper and glass slide. After discarding the empty glass slide, the collected biofilms were centrifuged at 4000 rpm for 10 minutes, and the supernatant liquid was carefully removed through pipetting. The resulting biofilm pellet was weighed and frozen at –20°C until subsequent DNA extraction. Biofilms were sampled right before adding the raw wastewater to assess the initial diversity after acclimatization of the microbiome (T0). Further, biofilms were destructively sampled on days 1, 4, 7 and 14 after the influent was added.

### DNA extraction

DNA extraction from biofilm and wastewater samples was carried out using the Qiagen DNeasy PowerSoil Pro Kit (Qiagen, Hilden, Germany) as per the manufacturers’ instruction. Sufficient quality and quantity of extracted DNA was assessed using a NanoDrop One spectrophotometer (Thermo Fisher Scientific, Waltham, MA, USA).

### High-throughput qPCR analysis for various ARGs, MGEs and genetic markers

To determine the relative abundance of target genes in the biofilm and wastewater samples, DNA extracts were sent to Resistomap Oy (Helsinki, Finland) for HT-qPCR analysis using a SmartChip Real-Time PCR system (TaKaRa Bio, Shiga, Japan). 10 ng DNA of each of the extracted samples were shipped to Resistomap for analysis. The target gene quantified based on previously published primers (Stedtfeld et al., 2018) included 27 ARGs and 3 MGEs along with the house keeping gene 16S rRNA gene for normalization of relative abundances (SI Table 1). Two fecal indicator taxonomic markers (*E. coli* & *Enterococcus*) were quantified simultaneously. The PCR reaction mixture (100 nL) was prepared using SmartChip TB Green Gene Expression Master Mix (TaKaRa Bio), nuclease-free PCR-grade water, 300 nM of each primer, and 2 ng/μL DNA template. After initial denaturation at 95 °C for 10 min, PCR comprised 40 cycles of 95 °C for 30 s and 60 °C for 30 s, followed by melting curve analysis for each primer set. Amplicons with non-specific melting curves or multiple peaks were excluded. The relative abundances of the detected gene to 16S rRNA gene were estimated using the ΔCT method based on mean CTs of three technical replicates (Fang et al., 2023). Any results with multiple peaks and amplification efficiency out of the range of 0.9 – 1.1 were discarded. The threshold cycle (Ct) of 31 was set as the limit of quantification (LOQ) The gene copies were calculated using Eq. 1 (Liu et al., 2018):

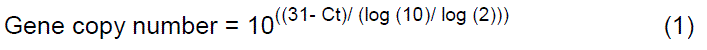

where Ct is the threshold cycle of the qPCR result, 31 is the LOQ-Ct, and log (10) / log (2) refers to the number of cycles of the ten-fold difference in the gene copy numbers, when the efficiency is 100%.

### Absolute bacterial abundance

To determine the absolute bacterial abundance on the biofilm carrier slides, the bacterial 16S rRNA marker gene was quantified using Real-Time qPCR in a C1000 Touch^TM^ Thermal cycler (Biorad, Hercules, CA, USA) and the 338F (CCTACGGGAGGCAGCAG) 518R (ATTACCGCGGCTGCTGG) primer set (Muyzer et al., 1993). As a qPCR standard the recombinant plasmid pBELX-1 (Bellanger et al., 2014) was extracted using the Wizard plus SV Miniprep DNA Purification System (Promega, Madison, WI, USA) according to the manufacturers’ instructions and linearized by restriction enzyme BamHI (Promega) before purification with the QIAquick PCR purification kit (Qiagen). Reactions were performed in technical triplicates in a MasterCycler RealPlex (Eppendorf, Hamburg, Germany) at a final volume of 20 μL with 10 μL of Luna® Universal qPCR Master Mix (New England Biolabs, Frankfurt, Germany). Each primer was added at a final concentration of 300 nM, and the reactions were run with 10 ng of DNA extract. The PCR program consisted of initial denaturation at 95 °C for 10 min and 40 cycles of denaturation (95°C; 15 s) and annealing and elongation (60°C; 1 min). Standard curves for the targets were created during every qPCR run, using the above-described standard plasmid with the standard target concentrations ranging from 10^6^ – 10^1^ copies per reaction. Standard curves with amplification efficiency 0.9 – 1.1 and R^2^ ≥ 0.99 were accepted and melting curve analysis was performed to assess the amplicons’ specificity. Screening for PCR inhibition was performed by spiking the standard plasmid into the DNA samples. No inhibition was detected in any of the samples. The limit of quantification was calculated for each individual qPCR run according to the MIQE guidelines (Bustin et al., 2009).

## 16S sequencing and analysis of microbial community diversity

The microbial community diversity and structure were profiled using high-throughput amplicon sequencing of the V3-V4 region of the 16S rRNA gene. DNA extracts were sent to the IKMB Kiel University (Germany), and the partial 16S rRNA genes were sequenced on an Illumina Novaseq using the primers (v3f: CCTACGGGAGGCAGCAG; v4r: GGACTACHVGGGTWTCTAAT;). Sequence analysis was carried out using Mothur v.1.37.1 (Schloss et al., 2009) according to the MiSeq SOP (Kozich et al., 2013) as assessed on 03.07.2022. Sequences were classified based on the RDP classifier (Wang et al., 2007). All sequencing data presented in this manuscript has been submitted to the NCBI sequencing read archive under accession number PRJNA1014398.

### Identification of invasion indicators

To investigate the invasion success of wastewater borne ARGs and bacteria, indicators were defined. Potential invasion indicators were defined as those targets from the HT-qPCR analysis that have a significantly higher relative abundance in wastewater compared to the biofilm samples at T0 and would hence be expected to rise upon exposure to the wastewater in the biofilms. Successful invasion indicators were then defined as the fraction of potential indicators that did indeed significantly increase in the exposed biofilms at T1 when compared to the control treatment. To determine significancy, a multilevel pattern analysis was carried out based on the relative gene abundances using the multipatt function of the indicspecies package in R (De Cáceres et al., 2010). For every mentioned computation the grouped biserial correlation and the phi correlation coefficients were calculated using a complex permutation design (n=9999) using the permute (Simpson et al., 2022) package in R. Using this setup, finally 6 of the 27 ARGs, 1 of the 3 MGEs and both fecal indicator taxonomic targets were defined as indicators of invasion success over time and used to determine temperature effects on invasion.

### Statistical analysis

All statistical analysis was performed in R Studio v4.1. (R core team, 2020). The relative abundance of each target gene derived from HT-qPCR was compared between control and exposed biofilms at different timepoints for each temperature group using a Wilcoxon rank-sum test. P-values were adjusted using the Benjamini & Hochberg method. Relative abundance below the detection limit (LOD) were set as zero before computation.

Pairwise comparison of the sum of abundance of all ARGs in the biofilms at T0 between different temperatures were tested using the student’s t-test. The change over time in the sum of relative abundance of all ARGs were tested for each treatment using the Kruskal-Wallis test. To test if the relative abundance of each individual ARGs across different temperatures in the control flumes follow a global trend with temperature the Wilcoxon rank-sum test was applied.

Trends regarding the relative abundance of target genes over time were tested based on Pearson correlation coefficients using the “ggplot” package (Wickham, 2016). To test if the slopes of the linear regression for the individual indicator genes over time follow a global trend based on temperature the Wilcoxon rank-sum test was applied.

Changes in bacterial diversity were assessed based on observed OTUs at 97% sequence similarity using the Bray-Curtis dissimilarity metric in the “vegan” package (Oksanen et al., 2022). Differences between the biofilm and wastewater community structure of control and exposed groups in different temperatures were tested using ADONIS in R v4.1.0 under the “vegan” package (Oksanen et al., 2022). Difference on the phylum level abundance between biofilm community at T0 and the wastewater community was tested using Kruskal-Wallis rank sum test.

Throughout the manuscript a p-value of p < 0.05 was determined as a statistically significant observed effect.

## Results

### Distinct river microbial biofilm communities based on incubation temperature

To study the effect of rising temperatures on AMR dynamics in rivers, river biofilms were grown on glass slides and set to acclimatize for one week at three climate chambers of temperature 20°C, 25°C, and 30°C. Thereafter, the biofilms for each temperature were subjected to two individual treatments: one group was exposed to wastewater, and a control group remained unexposed, with the experiment lasting for 14 days.

To determine if temperature on its own influenced the microbial community and resistome composition, biofilms from the control group were investigated first. At the initial timepoint (T0) after completing the one-week acclimatization period, river microbial communities clustered distinctly based on the incubation temperature (Fig. 1A). Replicate biofilms from flumes incubated at the same temperature clustered significantly together and apart from those incubated at different temperatures (p< 0.001, R^2^ = 0.37, ADONIS). Still, the most dominant phyla in the biofilms from all three temperatures were consistently *Proteobacteria* (57.87 ± 12.39% to 65.29 ± 7.50%), *Bacteroidetes* (8.27 ± 2.41% to 13.1 ± 2.19%) and *Firmicutes* (5.86 ± 6.75% to 1.59 ± 1.03%) (Fig. 1B). While a minor change in community composition was observed over time, the grouping based on temperature stayed consistent for the duration of the experiment (p< 0.001, R^2^ =0.29, ADONIS) (Fig. 1A). Furthermore, no statistically significant trend regarding the total amount of biomass based on 16S rRNA gene copies per cm² of the biofilm carrier was observed during the 14 days of recirculation in the flume. Bacterial numbers remained within the same order of magnitude throughout the experiment, suggesting that growth is a negligible factor in the experiment (Fig. 1C).

**Fig. 1:**
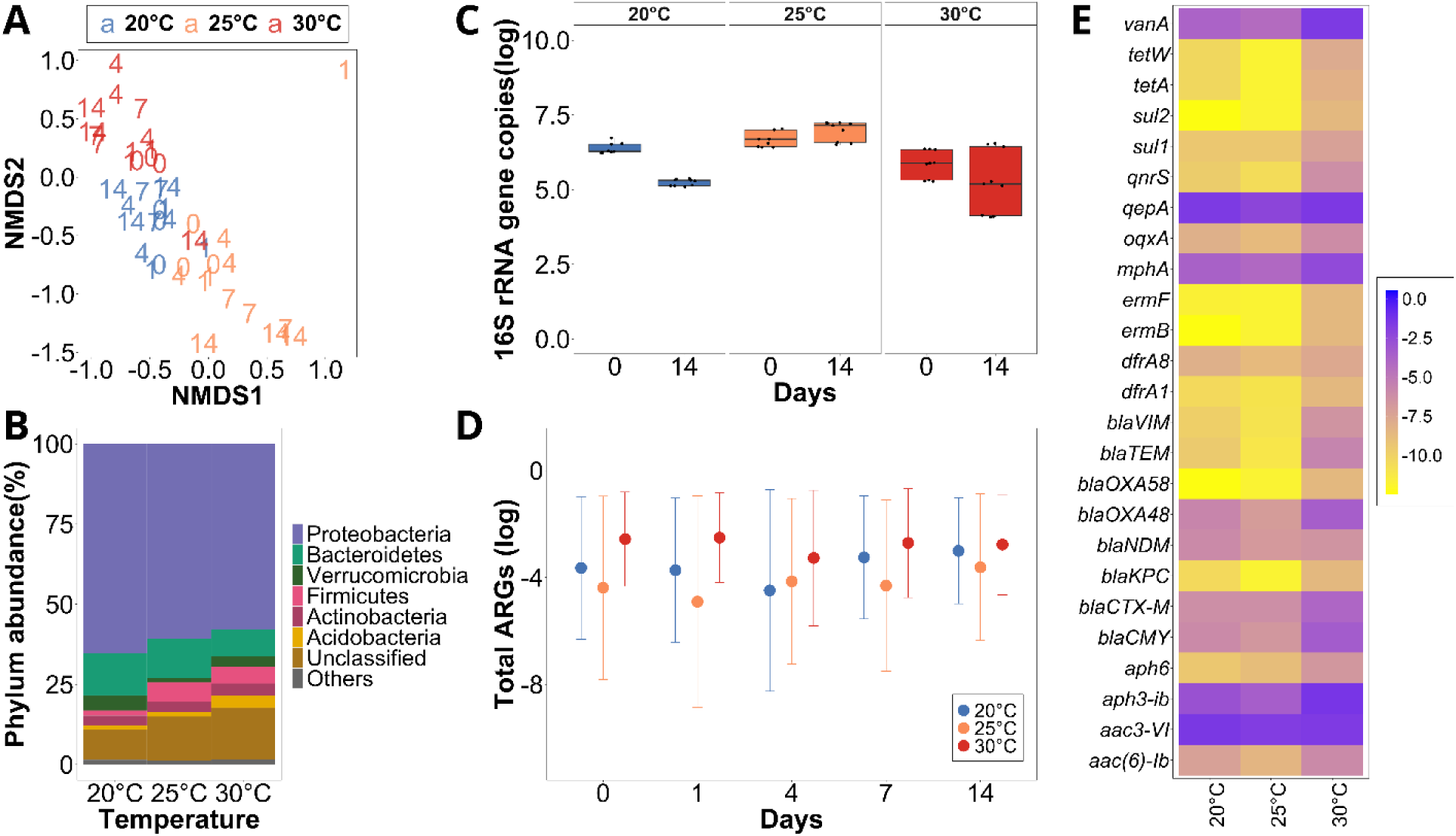
The initial diversity of the control biofilms after one week of acclimatization to the laboratory condition with different temperatures. A) NMDS (non-metric multidimensional scaling) plot using the Bray-Curtis dissimilarity to analyze the beta diversity of the biofilm for control group for T0, T1, T4, T7 and T14 for all the temperature groups. B) Relative phylum abundance in percent of the biofilms at T0 across all the temperature groups. C) Initial and final abundances of absolute 16S rRNA gene copies per cm^2^ in the biofilms across all temperature groups. D) Average of the sum of all tested ARG relative abundances normalized per 16S rRNA gene copies present in the biofilms for T0, T1, T4, T7 and T14 in the control group throughout the experiment across all the three temperatures. The error bars represent standard deviation. E) Heatmap showing the initial relative abundance of ARGs present in biofilms in control group for different temperatur groups.

### Characterization of the invading wastewater community

The wastewater microbial community was significantly dissimilar from the biofilm communities at T0 (p < 0.001, R^2^ = 0.29, ADONIS) (Fig. 2A). While the three dominant phyla, *Proteobacteria* (38.04 ± 27.8%), *Firmicutes* (33.95 ± 0.61%) and *Bacteroidetes* (13.26 ± 1.2%) were similar to the biofilms, significantly more *Firmicutes* and significantly less *Proteobacteria* were observed in the wastewater (Fig. 1B & 2B) (p < 0.05, Kruskal-Wallis rank sum test).

**Fig. 2:**
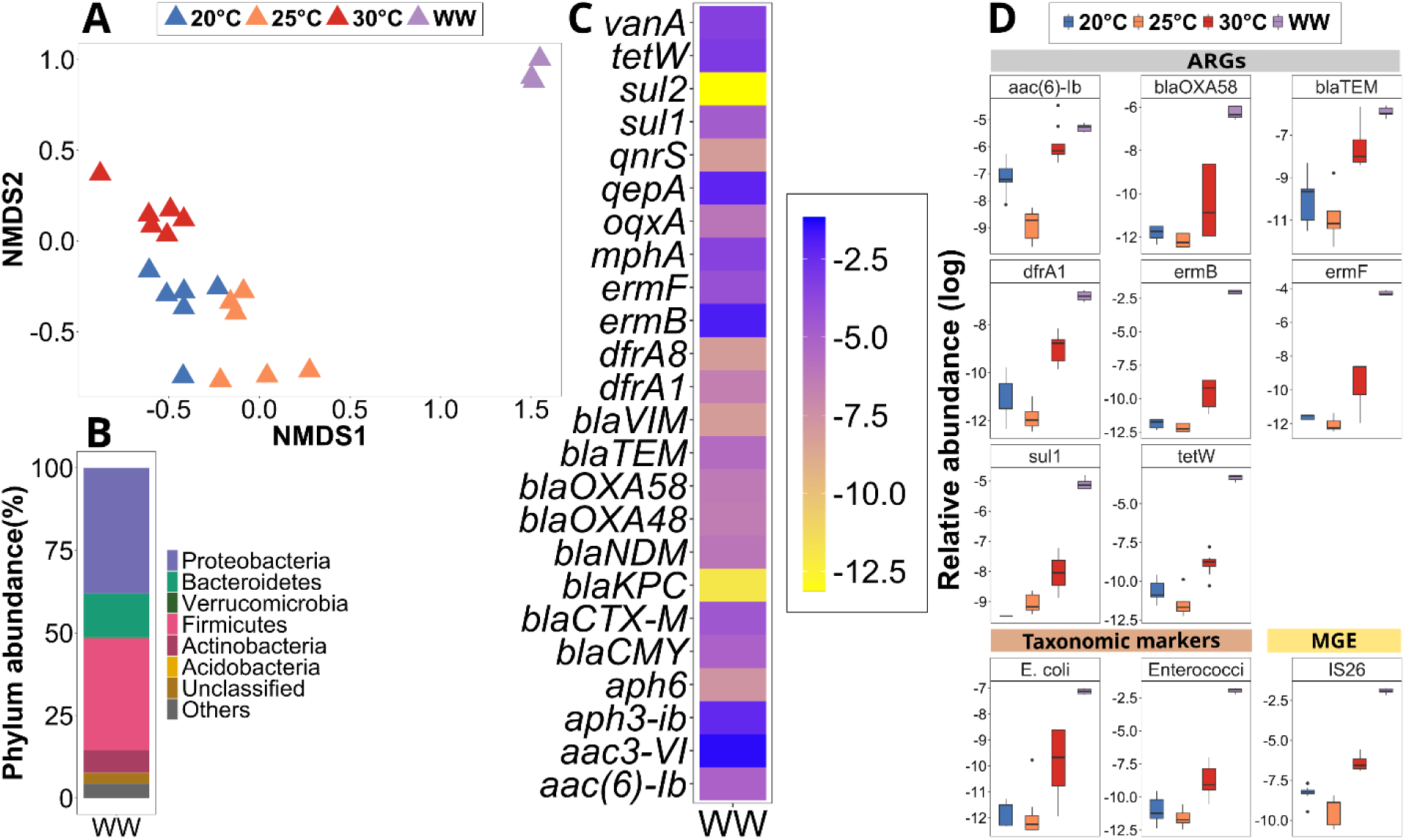
**Wastewater and biofilm composition (T0) and potential indicators of invasion before inoculation. A) NMDS (non-metric multidimensional scaling) plot using the Bray-Curtis dissimilarity to analyze the diversit of wastewater and the biofilm at T0 (stress = 0.1667) B) Relative phylum abundance in percentages of th wastewater. C) Relative abundance of all the ARGs tested for in wastewater before inoculation. D) Boxplot showing the initial relative abundances of ARGs, an MGE and two taxonomic markers identified as potential indicators of temperature groups at T0 and wastewater used for inoculation (n=9).**

### Elevated ARGs at higher temperatures in river biofilms

To determine if not only community composition, but also the resistome of the river biofilm communities were altered due to increasing temperatures, HT-qPCR of 27 ARGs was performed. The most abundant ARGs present in the control flume biofilms across temperatures were *aac3-VI* (0.15 ± 0.07 to 0.63 ± 0.19 ARG copies/16S copies), *qepA* (0.13 ± 0.05 to 0.35 ± 0.09), *aph3-ib* (0.08 ± 0.07 to 0.36 ± 0.34), *mphA* (0.024 ± 0.01 to 0.11 ± 0.02), and *vanA* (0.03 ± 0.02 to 0.17 ± 0.05) (Fig. 1E). These confer resistance to aminoglycosides, quinolones, macrolides and vancomycin. While the same set of ARGs were dominant across temperatures, the sum of all relative abundances of all the targeted ARGs was significantly higher in the biofilms exposed to temperature of 30°C with 1.82 ± 0.16 ARGs per 16S rRNA gene (Fig. 1E) compared to 20°C (0.60 ± 0.27, p < 0.01, paired t-test) and 25°C (0.28 ± 0.05, p < 0.01). However, there was no significant difference between the two lower temperatures (p = 0.354). This trend was mirrored at the individual ARG level where an increase in relative abundance with temperature was observed for a significant proportion of individual ARGs at 30°C compared to 20°C (p = 0.003) and 25°C (p < 0.001) based on a Wilcoxon rank-sum test. Similar to the microbial community composition, the relative abundance of ARGs for the control group did not undergo any significant changes over the 14 days of flume experiment (p > 0.2, Kruskal-Wallis test). However, the total ARG abundance at 30°C remained significantly elevated at all timepoints compared to 20°C and 25°C (p < 0.05, Kruskal-Wallis test) (Fig. 1D). This indicates that higher temperatures select for those community members that are hosting more ARGs initially, but that communities and their resistome are not affected by temperature once they reach a stable state after acclimatization.

### Identification of invasion indicator ARGs from wastewater into the river biofilm community

To quantify the invasion success of wastewater derived ARGs into the biofilm community, potential invasion indicators were defined as those targets from the HT-qPCR analysis that have a significantly higher relative abundance in wastewater compared to the biofilm samples at T0 and would hence be expected to rise in relative abundance in the biofilms upon exposure to the wastewater.

Among the 27 tested ARGs, the most abundant ones previously identified in the biofilms at T0 were also the most abundant in the wastewater: *aac3-VI* (0.25 ± 0.03 ARG copies/16S copies), *qepA* (0.1 ± 0.01), *aph3-ib* (0.08 ± 0.01), *mphA* (0.03 ± 0.003) _1and_ *vanA* (0.04 ± 0.004) (Fig. 2D).

O_t_her abundant ARGs in wastewater were *ermB* (0.13 ± 0.02), *tetW* (0.04 ± 0.006), and *ermF* (0.014 ± 0.001). These ARGs confer resistance to multiple antibiotic classes including aminoglycosides, quinolones, macrolides, vancomycin and tetracyclines, respectively.

Still, significant difference in relative abundance between wastewater and biofilms were observed: A total of 8 ARGs (*ermF*, *ermB*, *tetW*, *sul1*, *blaOXA58*, *dfrA1*, *blaTEM*, and *aac.6.Ib*) were identified as significantly more abundant in wastewater using multilevel pattern analysis (p < 0.01) (Fig. 2D) and selected as potential indicators of invasion. Additionally, two fecal bacterial indicators *E. coli* and *Enterococci* and one MGE (*IS26*), displayed a significantly higher association with the wastewater compared to the initial biofilms at all temperatures (p < 0.01) (Fig. 2D). An observable increase in relative abundance of these prevalent target genes in the biofilm upon exposure to wastewater would hence provide evidence successful invasion events into the river biofilms.

### Successful and temperature independent invasion of ARGs from the wastewater into the river biofilm community

To determine if invasion was indeed taking place, the abundance of the identified indicators in the river biofilms exposed to wastewater from all three temperature groups were tested one day after inoculation of the wastewater. Successful invasion indicators were then defined as the fraction of potential indicators that did indeed significantly increase in the exposed biofilms at T1 when compared to the control treatment. After one day, six of the eight ARGs (*blaOXA58, dfrA1, ermB, ermF, sul1 and tetW*) were significantly more abundant in the biofilm that was exposed to wastewater compared to those in the control flumes for each of the three different temperatures (all p < 0.05, Wilcoxon test) (Fig. 3). Especially, *ermB, ermF and blaOXA58*, which were below the detection limit in the majority of replicate biofilms from the control samples had high levels at T1 in the exposed biofilms indicating successful invasion from wastewater. After day 1 of invasion, for each of these six indicator ARGS, the relative abundance level was within one order of magnitude among different temperatures. Moreover, there was no clear trend observed with regards to the initial invasion efficiency across the different temperatures based on a Wilcoxon test.

**Fig. 3:**
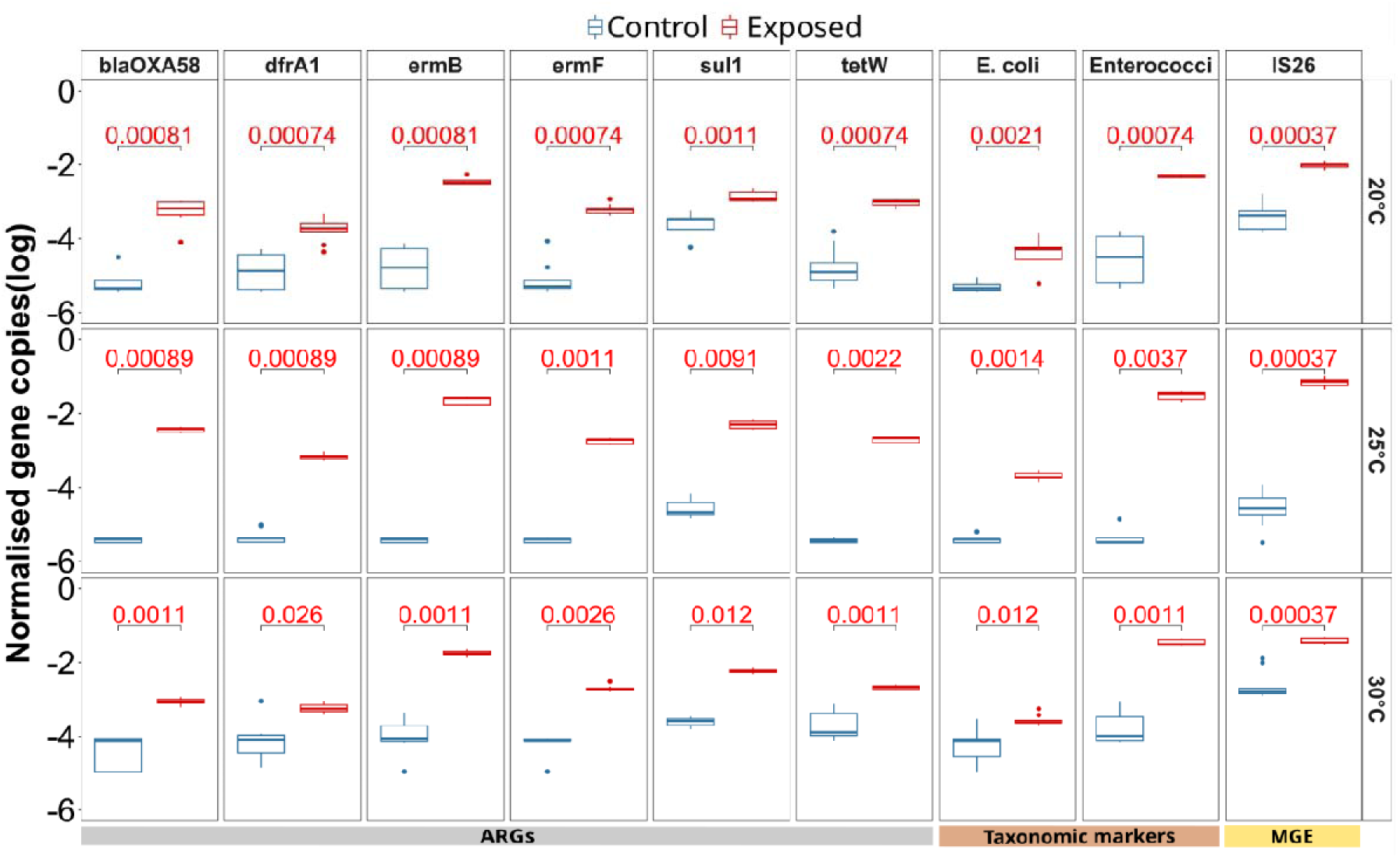
**Identification of indicator ARGs, taxonomic markers and MGEs for which a significant difference in relative abundance (log) was observed at day 1 after inoculation, compared between the wastewater exposed and the control biofilms across the three different temperatures (20°C, 25°C and 30°C). The p-value (Wilcoxon signed rank test) of the comparison for each gene (n=9), after applying Benjamin-Hochberg correction for multiple testing, is displayed at the top of each plot, with significant values (p < 0.05) displayed in red.**

Notably, the remaining 2 ARGs significantly associated with wastewater (*blaTEM*, and *aac.6.Ib*) did not significantly increase in abundance across all temperatures (Fig. S2), indicating that the invasion of these indicator ARGs is not merely stochastic but potentially connected to the specific ARG hosts that might have different invasion efficiency into the biofilm microbiome. Importantly, for the remaining 19 ARGs where the relative abundance in the wastewater and the T0 biofilm communities displayed no significant difference and that were hence previously excluded as potential indicators of invasion, no consistent significant increase across all temperatures was observed (p > 0.05 for at least one temperature per ARG, Wilcoxon test; Fig. S2). Among the three potential non-ARG indicators, the MGE *IS26* (p < 0.001 Wilcoxon rank sum test), *E. coli* (p < 0.05, Wilcoxon rank sum test) and *Enterococci* (p < 0.001, Wilcoxon rank sum test) showed a significant increase on day 1 after inoculation for all three temperatures (Fig. 3). In summary, a clear indication of successful initial invasion by ARGs or ARBs from wastewater into the biofilms independent of temperature was detected right after the introduction on day 1, which allows investigating the fate of these invaders from wastewater over time as a function of temperature.

### Invasion indicator genes and organisms are lost more rapidly with increasing temperature

To understand the fate of the invaders over time after initially entering the biofilms, biofilms were sampled at days 1, 4, 7 and 14. At the highest temperature (30°C) for all nine indicators (6 ARGs, 1MGE, *E. coli*, *Enterococci*) a clear decline in abundance with a significantly negative slope based on Pearson correlation was observed (all p < 0.05; Fig. 4). While similarly declines in abundance over time were observed for the majority of indicators at 20°C and 25°C, the slopes were less steep, demonstrating a lower loss rate of indicators over time (Fig. 4). This manifested statistically, as overall, the loss rate of each of the invading nine indicator genes and organisms over time was significantly higher at 30°C compared to 20°C and 25°C (both p < 0.01) based on a Wilcoxon signed-rank test for temperature, while no significant difference in loss dynamics was observed between the two lower temperatures. In certain cases, such as *blaOXA58* (p = 0.67, Pearson Correlation), and *sul1* (p = 0.50) at 20°C there was no significant decrease in abundance over time observed for the invading ARGs, and their abundance remained stagnant. The genes *sul1* and *ermF* even slightly but significantly increased over time at 25°C (both p < 0.05). Contrary to the wastewater inoculated flumes, not a single significant decrease over time was detected in the control treatment for either of the indicator genes at any temperature (all p > 0.05; Fig. SI 3).

**Fig. 4:**
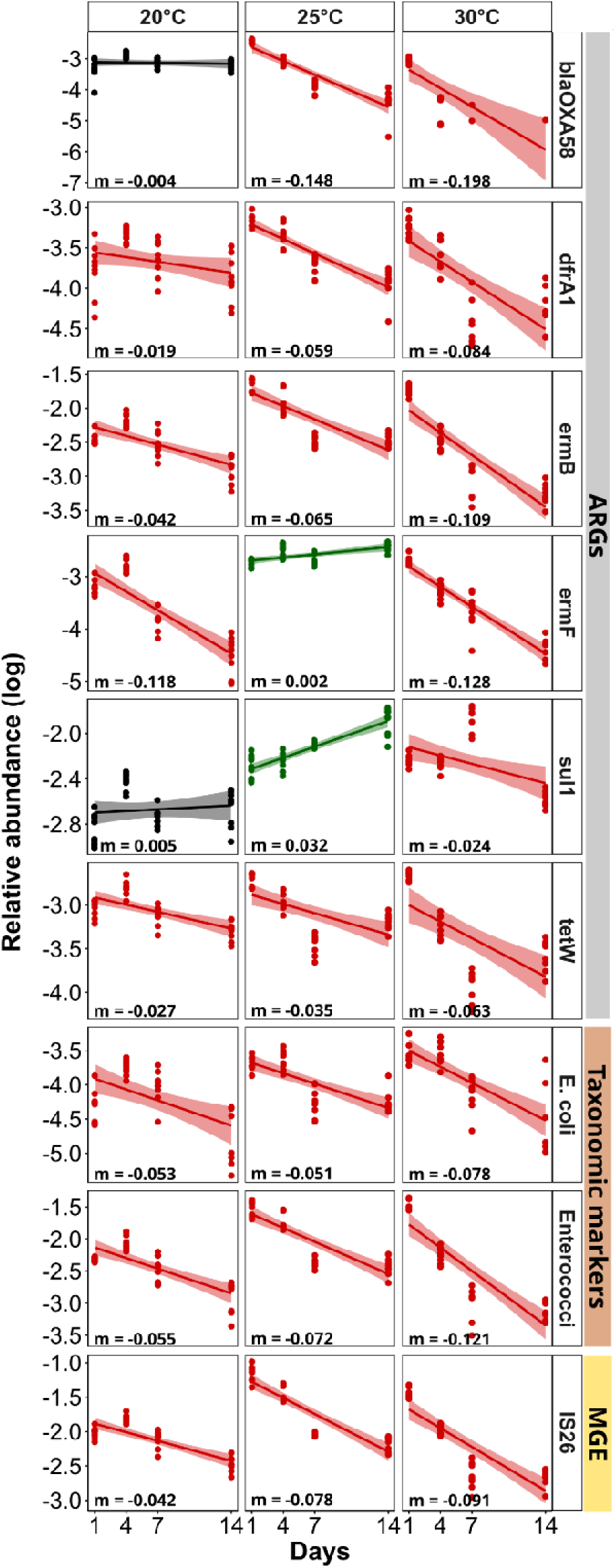
**Temporal dynamics of the nine identified indicator genes in wastewater invaded biofilms across three temperature groups. Linear regression for 16SrRNA gene normalized relative indicator gene abundance (log transformed). Red lines indicate a significant decrease, black lines no significant change, green lines a significant increase of the marker gene over time based on Pearson correlation analysis (p < 0.05).**

The temperature dependent loss dynamics were also reflected in the outcome of the flume experiment. On day 14 at 30°C, the relative abundance of five out of the six invading ARGs and all three non-ARGs, the MGE IS*26* and the two fecal indicator bacteria *E. coli* and *Enterococci* had returned to similar abundance level observed in the control flume. Only the indicator ARG *sul1* remained significantly elevated compared to the control group (p < 0.05, Wilcoxon test) (Fig. 5). Contrary, at 20°C, three of the indicator ARGs (*blaOXA58, ermB* & *tetW*), and at 25°C, all indicator ARGs remained significantly elevated (p < 0.01, Wilcoxon test). Similarly, the three non-ARG indicators remained at higher levels than those observed in the control at the two lower temperatures on day 14 (p < 0.05, Wilcoxon test).

**Fig. 5:**
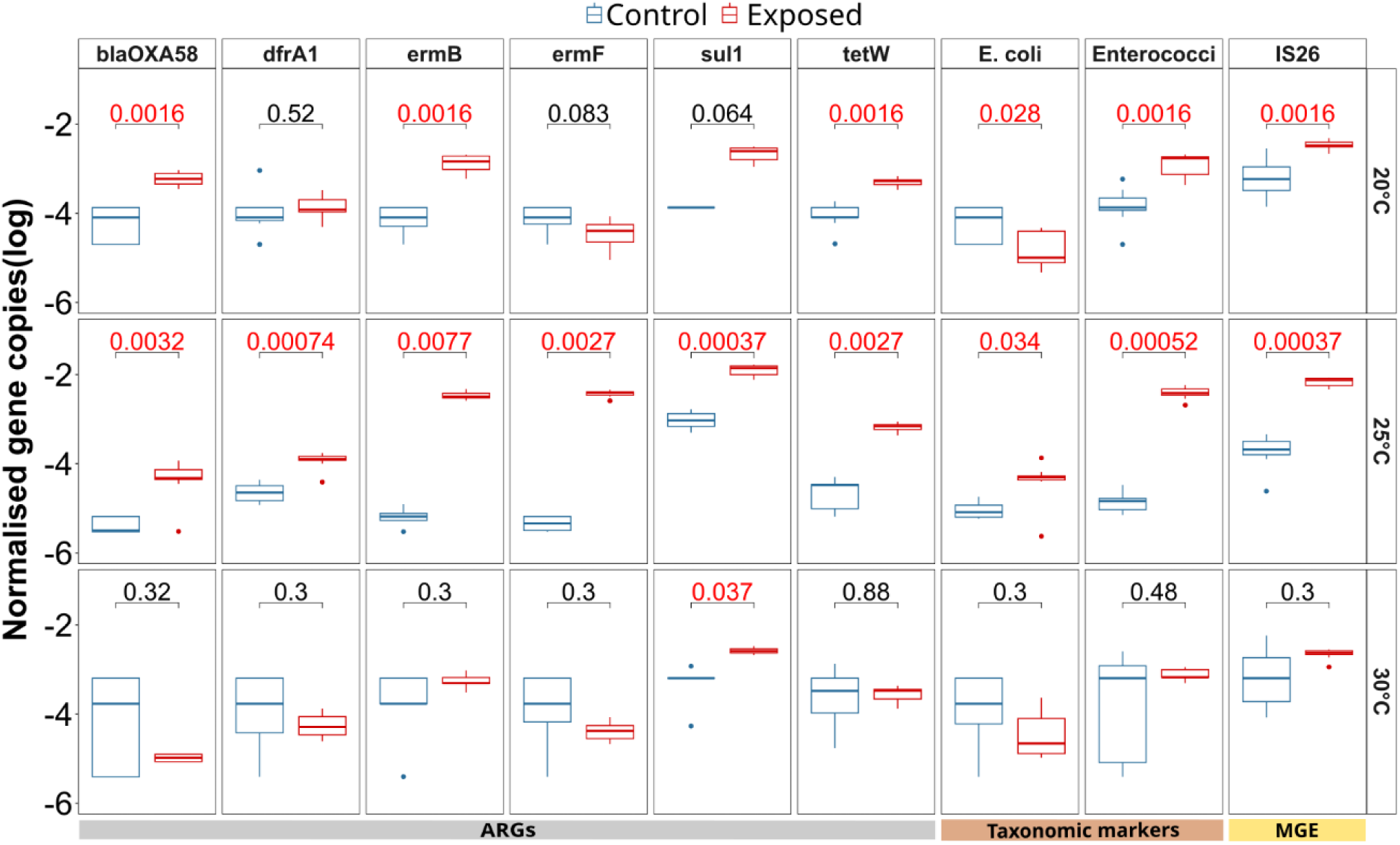
**Indicator ARGs, taxonomic markers and MGEs relative abundance in log scale at day 14 compared between the wastewater invaded and the control biofilms across the three different temperatures (20°C, 25°C and 30°C). The p-value (Wilcoxon test) of the comparison for each gene after correction for multiple testing i displayed at the top of each plot, with significant values (p < 0.05) displayed in red.**

Overall, this suggests that the loss of invading ARGs and organisms at 30°C is not only rapid but that the river biofilm is resilient against these invaders and can return to its original level before invasion. Contrary, at lower temperatures effects of the invasion event remain detectable for an elevated amount of time. Importantly, similar to the control flumes, no change in biomass was detected over the course of the experiment at either temperature again indicating that growth is a negligible factor for the observed temperature dependent effects.

### Higher resilience of the microbial community composition with increasing temperature

To determine if the observed initial invasion into the biofilm with subsequent loss of invaders is not only reflected on the genetic, but also on the community composition level, the microbial diversity dynamics over time of the biofilms were explored. For neither the controls nor the wastewater invaded river biofilms major changes in the phylum level distribution over time were observed, with Proteobacteria, Bacteroidetes and Firmicutes remaining the main observed phyla throughout (Fig. S4). However, on the beta diversity level based on OUT abundances the wastewater invaded biofilms displayed major changes over time (Fig. 6). Throughout, all biofilm samples cluster significantly apart from the initial invading wastewater community (p < 0.05, R^2^ =0.294, ADONIS). Initially, the control and influent biofilms before treatment clusters together for all temperatures and only minor changes in beta diversity over time were observed for the control biofilms. However, on day 1 after the wastewater community was added to the flume, biofilms from the wastewater group clustered immediately between the initial biofilm as well as the initial wastewater samples but remained statistically different in diversity from either of the initial microbiomes (p < 0.05, R^2^ = 0.074, ADONIS). This trend was apparent across all three temperatures, suggesting indeed wastewater bacteria became part of the biofilm community and affected its diversity throughout (Fig. 6). At the two lower temperatures (20°C & 25°C), the wastewater invaded biofilms, while significantly shifting in diversity, remained distinct and dissimilar to the initial biofilm, the control biofilms as well as the initial wastewater community. Contrary, the wastewater invaded biofilms at 30°C rapidly return to cluster with the control biofilms, with the two treatment groups showing no significant distinction after 14 days when considering both, treatment and day in the ADONIS test (p = 0.12) (Fig. 6). Hence, the previously detected resilience at the ARG level at the higher temperature is also realized at the microbiome level, with microbial diversity and composition returning rapidly back to the initial state after the invasion event.

**Fig. 6:**
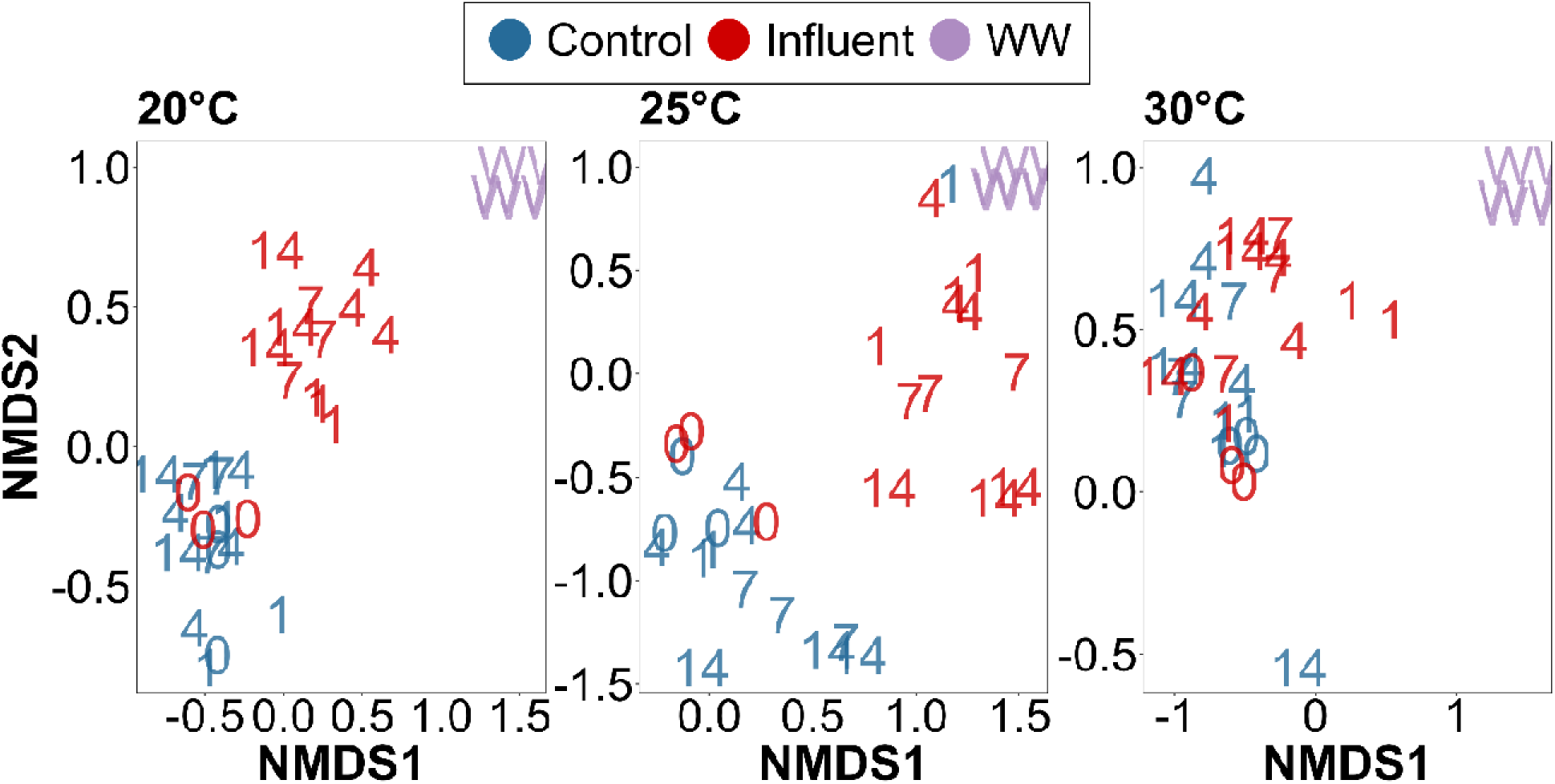
**Non-metric multidimensional scaling (NMDS) plot for beta-diversity of the river biofilm communities calculated using the Bray-Curtis dissimilarity. The NMDS plot for three temperature groups Tm20, Tm25, Tm30, and WW influent showing structural variations of the microbial community composition between the control and the treatment group.**

## Discussion

In this study we demonstrate how increasing temperatures could affect the dynamics of antimicrobial resistance in river biofilm communities. We observe tradeoffs between an increase in the pre-existing AMR present in the biofilm community and a decrease in invadability of the biofilms with novel ARGs at elevated temperatures. Consequently, when estimating effects of global warming on environmental resistomes both these mechanisms need to be taken into account. Understanding such ecological mechanisms of AMR proliferation is especially relevant for interpreting environmental data when setting up environmental AMR monitoring frameworks (Bengtsson-Palme et al., 2023; Klümper et al., 2022).

To determine effects of warming on resistome dynamics, biofilms were grown on glass slides from a German river for several weeks, retrieved and placed into laboratory flume recirculation systems at different elevated temperatures. As the sampling location is close to the spring of the river, and no wastewater or agricultural runoff is entering upstream, the biofilms can be considered as relatively low anthropogenically impacted (Klümper et al., 2023). Hence, any observed ARGs are likely part of the natural river resistome. Already after one week of acclimatization at the respective temperatures, biofilms in control flumes exposed to 30°C displayed a significantly higher abundance of overall as well as individual ARGs when compared to those exposed to 20°C and 25°C. This difference in ARG abundance remained throughout the experimental observation period of 14 days, however, no further increases with time were observed. In addition to the observed increases in resistance, communities from the different temperature treatments were significantly different. Similar trends of ARGs increasing with temperature were equally detected in a long-term soil warming trial (Li et al., 2022) and equally attributed to shifts in the microbial community.

From an ecological perspective the observed dynamics can be explained based on the ARG hosts intrinsic fitness and/or the fitness of being resistant itself. First, if those bacteria hosting ARGs have a higher intrinsic fitness at higher temperatures they would likely make up a higher relative proportion of the biofilm at elevated temperatures (Rodríguez-Verdugo et al., 2020). In addition, the fitness cost of hosting ARGs is often temperature dependent (Rodríguez-Verdugo et al., 2020, 2013; Trindade et al., 2012), with these studies providing examples of how resistance is not necessarily costly or selected against under thermal stress in the absence of any selection pressure through antibiotics. However, equally increases in the costs of resistance at higher temperatures have been reported (Gifford et al., 2016). Our study suggests that the former, lower costs of resistance at higher temperatures is likely to play a role in river biofilms, hence leading to the observed increase in ARGs.

While our study addresses increases in AMR with warming from an environmental, whole-community perspective, the observed effects were mirrored in resistance profiles of specific pathogens on a clinical level (Li et al., 2022; McGough et al., 2020): Based on an ecological analysis of AMR prevalence in 4Lmillion tested pathogens across 28 European countries between 2000 and 2016, countries with warmer ambient minimum temperatures, experienced more rapid resistance increases ranging from 0.33-1.2% per year across all antibiotic (McGough et al., 2020). Further, in a study across 28 Chinese provinces between 2005 and 2019, a 1°C increase in average ambient temperature was associated with a 6-14% increase in carbapenem resistance in *K. pneumoniae* and *P. aeruginosa* (Li et al., 2022). These effects held true, even after accounting for known drivers of resistance such as antibiotic consumption and population density.

Aside from temperature effects on the natural resistome of river biofilm communities we studied the invasion dynamics of ARGs from a wastewater community into these river biofilms at the three different temperatures (20, 25 & 30°C). Successful initial invasion of the incoming resistant bacteria and ARGs into the biofilm was observed shortly after their introduction to the flumes at day 1. Specifically, six ARGs, one MGE and two taxonomic markers were detected at higher relative abundance than in the control biofilms, with relative abundances increasing by several orders of magnitude across all temperatures. The previously discussed increases in ARGs due to temperature were relatively minor to those observed due to invasion from wastewater which were orders of magnitudes higher. Among the nine invasion indicator markers, the ARGs are well known to be associated with anthropogenic activities and WWTPs (Karkman et al., 2019; Pruden et al., 2006; Zhang et al., 2022) and were identified in our wastewater sample at far elevated abundance compared to the original biofilms. The taxonomic markers, *E. coli* and *Enterococci,* are well-known fecal pollution indicators (Garner et al., 2017; Karkman et al., 2019) and multi-drug resistance has been regularly reported for both (De Oliveira et al., 2020). Therefore, *E.coli* and *Enterococci* can be considered as model invading resistant bacteria for this study.

With regards to temperature, on day 1 no clear distinct pattern regarding the initial invasion success of the nine invasion indicators was observed between the temperature groups. This suggests that initial introduction of the invaders to the biofilm mainly depends on the propagule pressure, hence the number of invading bacteria or genes (Acosta et al., 2015; Kinnunen et al., 2018). This coincides with the result from a previous study using the same setup where the biofilms were invaded by *E.coli* in presence and absence of copper induced stress, where the initial invasion success of *E.coli* entering the biofilm was purely stochastic and independent of stress (Bagra et al., 2023). Furthermore, the successful initial invasion was equally observed in the community composition of the biofilms. Day 1 biofilm communities consistently grouped between the original biofilm and the wastewater samples, confirming that community coalescence in the biofilm as a result of initial invasion is indeed taking place.

Over time after the initial invasion event, a decline in the relative gene copy number of the invasion indicators was observed for all the temperature groups. However, this decline was far more rapid and pronounced for all markers at the highest temperature (30°C). This resulted in indicator levels at 30°C returning to similar levels than those of the non-invaded control biofilms after 14 days. On the contrary, the majority of indicators remained significantly elevated compared to the control biofilm at 20°C and 25°C. Considering that the natural levels of ARGs in the 30°C flumes were significantly elevated, as discussed earlier, it is important to note that the levels observed in the invaded 20°C and 25°C biofilms remained above the levels obtained in the 30°C control flume. Similar dynamics were observed with regards to community composition where no difference between the control and the invaded biofilms were observed at 30°C, while at 20°C and 25°C the communities of the biofilms remained significantly different. This is pointing towards an increased resilience at higher temperatures of the river biofilm.

When considering such invasion events within the framework of community coalescence, it can be expected that invading species replace indigenous species, if they are better adapted to the prevailing environmental conditions (Castledine et al., 2020; Rillig et al., 2015; Rocca et al., 2020). Considering that many of the resistant wastewater bacteria are of fecal origin their temperature optima are likely to be higher than those of the indigenous river biofilm community. For example, for our two model invader species the temperature optimum of *E. coli* is 37°C to 42°C (Gonthier et al., 2001) and for *Enterococci* is 42°C to 45°C (Fisher and Phillips, 2009), which would suggest that their invasion success should be higher at elevated temperatures.

However, higher temperatures also increase the metabolic rates of the indigenous community members, hence leading to a faster depletion of resources, resulting in a higher level of competition through limited resource access towards the invading species. This increased competition, together with the fact that many fecal bacteria struggle to display significant growth rates under environmental nutrient conditions independent of temperature (Anjum et al., 2021; van Elsas et al., 2011), could lead to the observed faster elimination of invading bacteria and their resistance genes. In addition, priority effects, which regularly determine the outcome of coalescence events (Fukami et al., 2007), could here be at play if the indigenous biofilm is already established before invasion, and could play an increased role when the biofilm bacteria show an increased level of activity at higher temperatures. While short term change in temperature could lead to an altered and less stable microbial community structure (Radujković et al., 2018), here biofilms were left to acclimatize to the temperature for one week before inoculation and remained stable in composition thereafter. This suggests that the community adapted to the respective temperatures and can be considered as established, meaning priority effects including an inherent advantage against invading bacteria are at play. Together, these factors contribute to the observed dynamics, where a higher resilience of the river biofilm community towards invasion by resistant bacteria and their ARGs are observed at 30°C. Similar effects on the temperature dependence of coalescence in aquatic bacterial communities were observed when mixing marine communities across different temperatures, where resistance of aquatic bacterial communities to invasion was shown to be strengthened by warming (Vass et al., 2021). This allows to presume that the here observed effects are likely to be transferable to other scenarios where invasion takes place.

Notably, the here described temperature effects did not follow a linear trend with increasing temperature. While consistently significant differences between the 30°C and the two lower temperature treatments were observed, the biofilms at 20°C and 25°C behaved rather similarly with regards to levels of natural resistance in control biofilms as well as invasion dynamics of foreign ARGs from wastewater. Temperature dependencies of microbial processes or ecosystem services regularly do not follow linear trends, as the influence of temperature on them is mediated by the temperature sensitivity of the individuals making up the microbial community and their biotic interactions (Johnston and Sibly, 2018; Yvon-Durocher et al., 2012). Rather temperature thresholds are crossed at which the temperature dependence switches between low, intermediate and high temperatures, as has for example been demonstrated for global ecosystem respiration rates (Johnston et al., 2021). Our results suggest that temperature effects regarding the spread of resistance in river microbial communities equally follow rather a threshold than a linear temperature model.

Overall, we here demonstrate tradeoffs of increasing temperature on the dynamics of AMR in river biofilm communities. While an increase in natural resistance in the biofilm at elevated temperatures was apparent, it coincided with a decrease in invadability by foreign ARGs originating from wastewater. This implies that estimating effects of climate change on river microbial communities might highly depend on which of the two processes dominates in the specific river ecosystem.

## Acknowledgements

The authors thank Christiane Zschornack and Steffen Kunze for help in sampling, setting up and running the flume experiments and molecular analysis. We further thank the members of the ANTIVERSA consortium for their contributions in designing the flume setup.

## Funding

This work was supported by the ANTIVERSA project funded by the Bundesministerium für Bildung, und Forschung of Germany under grant number 01LC1904A. KB was supported through a DAAD scholarship in the program Research Grants – Bi-nationally Supervised Doctoral Degrees/Cotutelle, 2021/22 (57552338). UK & TUB were supported by the Urban Resistome project funded by the Deutsche Forschungsgemeinschaft (DFG) under project number 460816351 and the PRESAGE project funded by the Bundesministerium für Bildung und Forschung under grant number 02WAP1619. GS was supported through a Faculty Initiation Grant (FIG/100762) and the Indian Department of Science and Technology under grant number DST/TM/INDO-UK/2K17/46. CC, AT-B, and ES were supported by the ERANET-BIODIVHEALT-ANTIVERSA project within PNCDI III of the Romanian National Authority for Scientific Research and Innovation under grant number 117/2020. AT-B and ES were also partially supported through the Core Project BIORESGREEN, subproject BioClimpact number 7/30.12.2022. Responsibility for the information and views expressed in the manuscript lies entirely with the authors.

## CRediT authorship contribution statement

Kenyum Bagra: Conceptualization; Investigation; Formal analysis; Validation; Visualization; Data Curation; Methodology; Funding acquisition; Writing – Original Draft; Writing – Review & Editing.

David Kneis: Formal analysis; Visualization; Supervision; Writing – Review & Editing. Daniel Padfield: Visualization; Writing – Review & Editing.

Edina Szekeres: Methodology; Resources; Writing – Review & Editing. Adela Teban-Man: Methodology; Resources; Writing – Review & Editing.

Cristian Coman: Methodology; Resources; Supervision; Funding acquisition; Writing – Review & Editing.

Gargi Singh: Conceptualization; Supervision; Writing – Review & Editing; Thomas U. Berendonk: Conceptualization; Methodology; Resources; Supervision; Project administration; Funding acquisition; Writing – Review & Editing;

Uli Klümper: Conceptualization; Formal analysis; Validation; Visualization; Data Curation; Methodology; Resources; Supervision; Project administration; Funding acquisition; Writing – Original Draft; Writing – Review & Editing.

## Competing Interests

The authors declare no competing interests.

## Data and materials availability

The main datasets supporting the conclusions of this article are included within the article. Original sequencing data is available in the NCBI sequencing read archive under project accession number PRJNA1014398. Any additional data is available through the corresponding author upon reasonable request.

## Notes

### Competing Interest Statement

The authors have declared no competing interest.

