## Supplementary Information for "Tradeoffs of increasing temperatures for the spread of antimicrobial resistance in river biofilms"

01062 Dresden, Zellescher Weg 40,

Germany

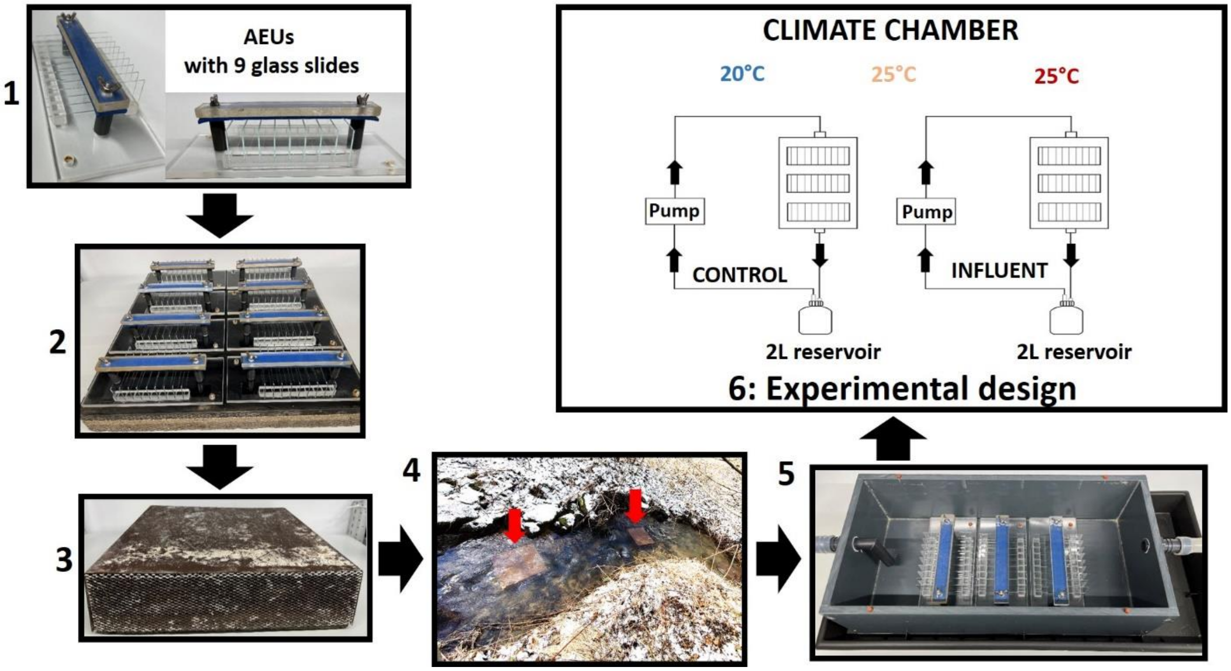


**SI Fig. 1: Schematic workflow of the flume experiments adapted from** (Bagra et al., 2023)**. 1) Biofilms were grown on microscope glass slides of 76 x 26 mm size, with 9 glass slides each fixed in artificial exposure units (AEUs) constructed from Plexiglas. 2) 8 AEUs were screwed onto a concrete plate and 3) covered with a stainless-steel cover providing shade and protection with a metal mesh at the front and the back allowing river water to stream through the AEUs. 4) The AEUs were immerged in the river Hirschbach (HIR) for one month to allow biofilm growth. 5) The AEUs containing the glass slides with grown biofilms were then transferred to laboratory artificial flume systems (40 cm long, 19.1 cm width, 10 cm height), made of polypropylene, and equipped with an in- and an outflow pipe at either end. Each flume was connected to a 2L reservoir connected to a recirculation pump and filled with filter sterilized river water constantly recirculated with a pump. 6) Three replicate AEUs containing glass slides covered with river biofilms were placed in the middle of each flume and exposed to three different temperatures for a period of 7 days acclimatization. Thereafter 2 different treatments were applied: a) 20% volume of the flume river water was replaced with wastewater**(Mahoney et al., 2021) **influent and b) a control group was run without exposure to wastewater. Glass slides containing biofilms were then destructively sampled to monitor the invasion success of wastewater specific resistance genes and bacteria into the biofilm communities.**

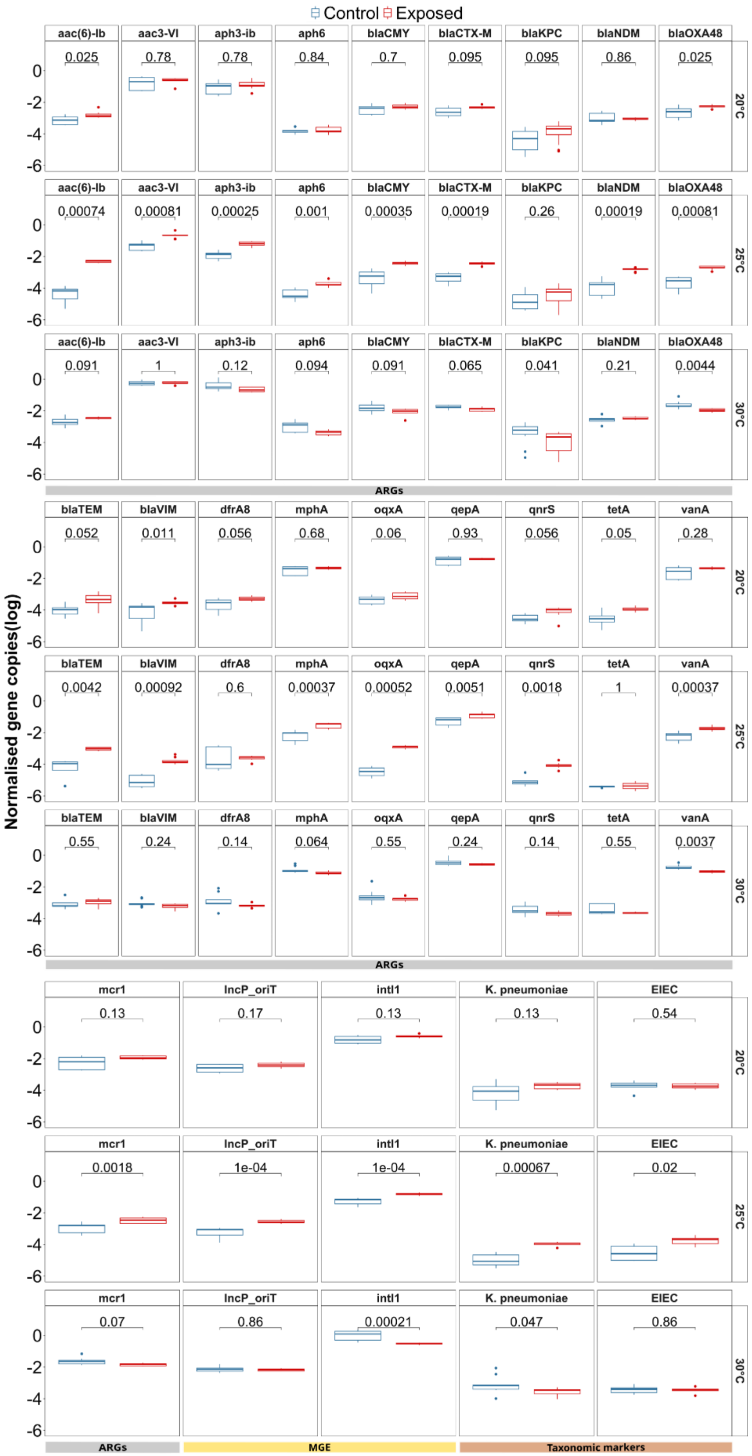


**SI Fig. 2: : ARGs, taxonomic markers and MGEs for which no significant difference in relative abundance (log) across all temperatures was observed at day 1 after inoculation, compared between the wastewater exposed and the control biofilms across the three different temperatures (20°C, 25°C and 30°C). The p-value (Wilcoxon signed rank test) of the comparison for each gene (n=9), after applying Benjamin-Hochberg correction for multiple testing, is displayed at the top of each plot.**

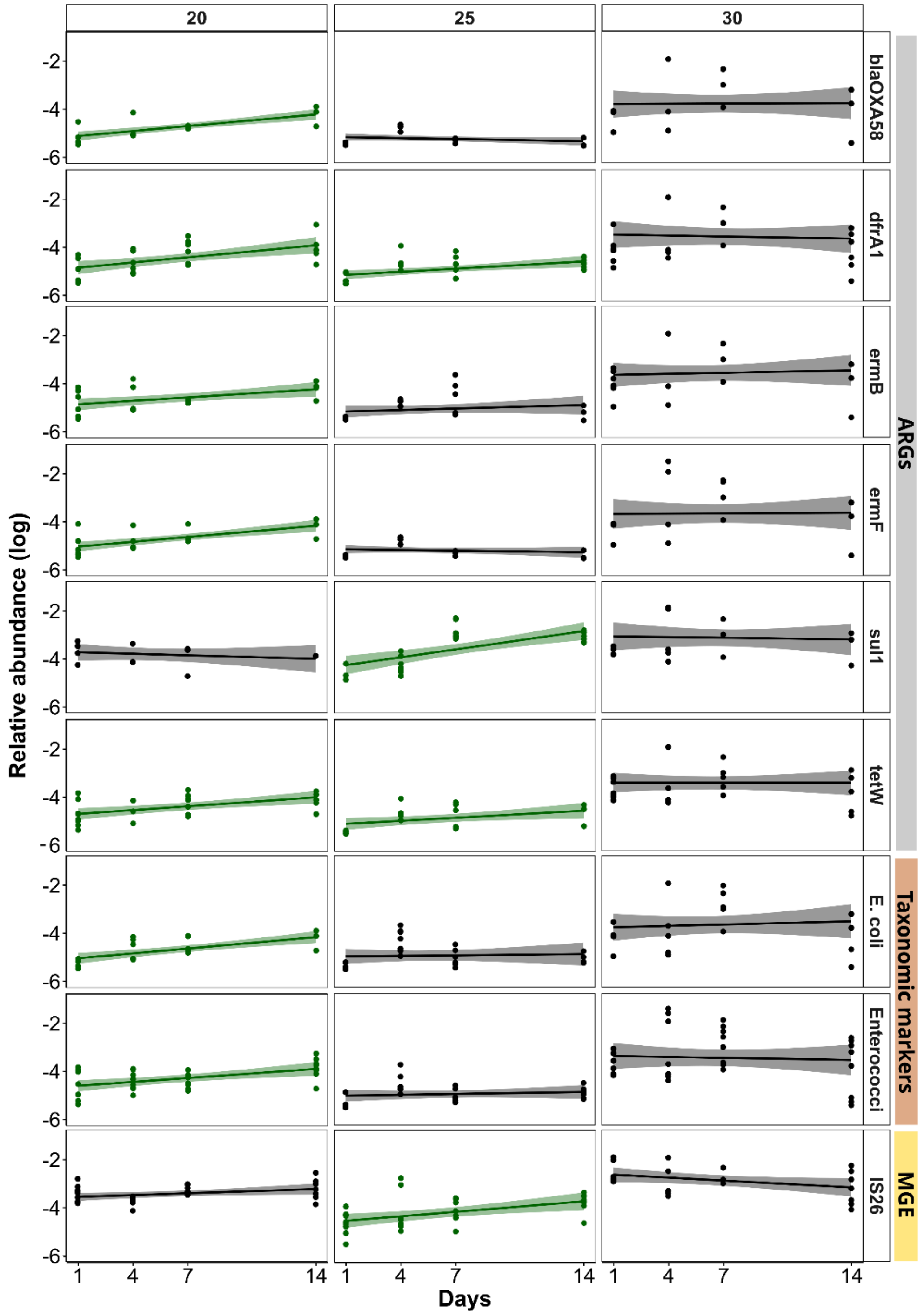


**SI Fig. 3: Temporal dynamics of the nine identified indicator genes in control flumes across three temperature groups. Linear regression for 16SrRNA gene normalized relative indicator gene abundance (log transformed). Red lines indicate a significant decrease, black lines no significant change, green lines a significant increase of the marker gene over time based on Pearson correlation analysis ($p < 0.05$).**

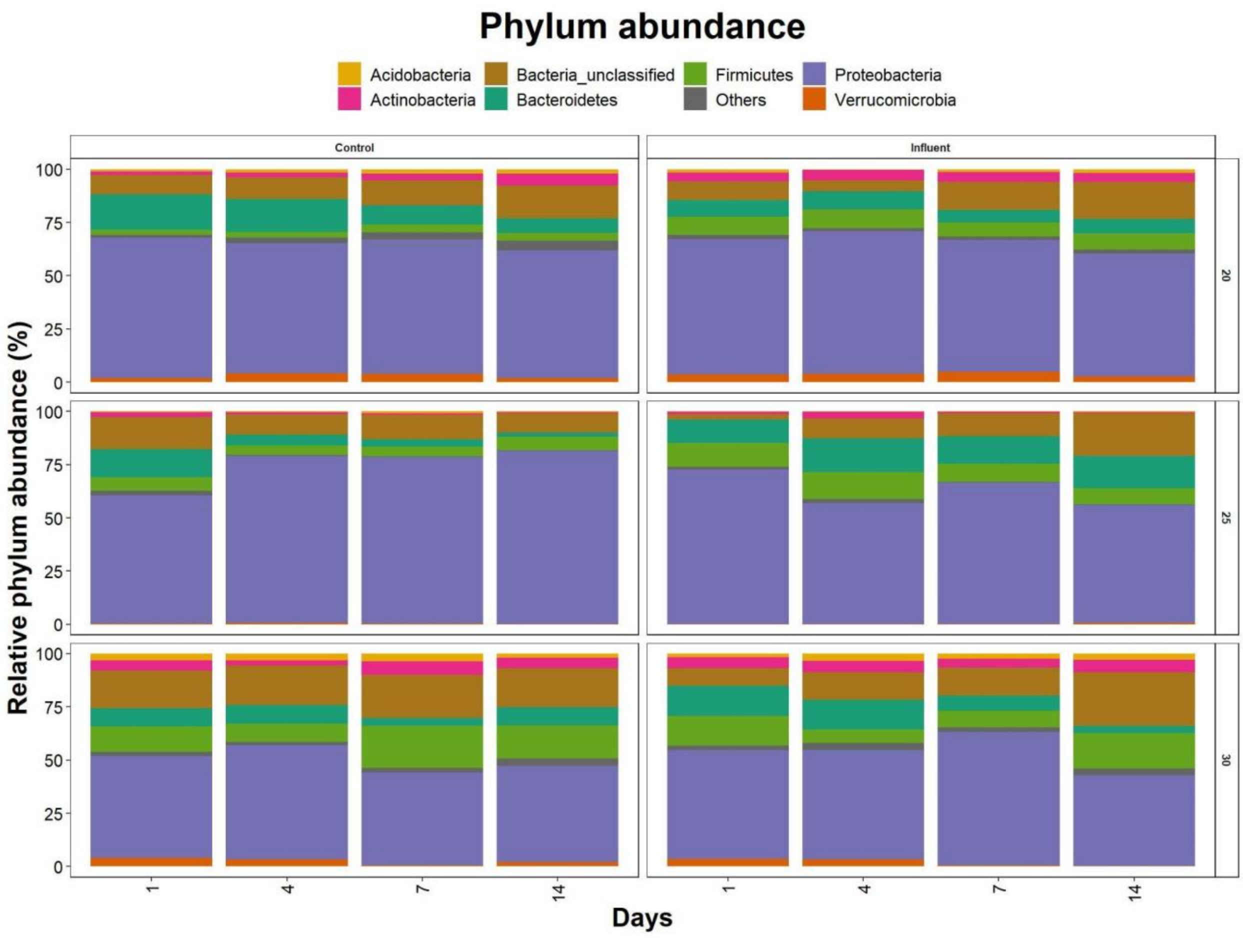


**SI Fig. 4: Microbial community composition at the phylum level over time in biofilms of control and influent exposed treatment across temperatures.**

**SI Table 1: Monitored indicator ARGs, MGEs and bacteria using HT-qPCR including primers used for amplification**

| Primer name | Gene/ Organism | Target antibiotic class/ Target type | Forward Primer | Reverse Primer |
|---|---|---|---|---|
| 16S rRNA | 16S rRNA | Bacterial abundance | GGGTTGCGCTCGTTGC | ATGGYTGTCGTCAGCTCGTG |
| aph3-ib | *aph3-ib* | Aminoglycoside | AACAGGTTTGGGAGGCGATG | CGCAACAAGCCTCTCCTGAA |
| aac3-VI | *aac3-VI* | Aminoglycoside | CGTCACTTATTCGATGCCCTTAC | GTCGGGCGCGGCATA |
| aph6 | *aph6* | Aminoglycoside | CCCATCCCATGTGTAAGGAAA | GCCACCGCTTCTGCTGTAC |
| aac(6')-Ib_2 | *aac(6')-Ib* | Aminoglycoside | GTTTGAGAGGCAAGGTACCGTAA | GAATGCCTGGCGTGTTTGA |
| blaVIM | *blaVIM* | Beta Lactam | GCACTTCTCGCGGAGATTG | CGACGGTGATGCGTACGTT |
| blaNDM | *blaNDM* | Beta Lactam | GGCCACACCAGTGACAATATCA | CAGGCAGCCACCAAAAGC |
| blaCMY_2 | *blaCMY* | Beta Lactam | AAAGCCTCAT GGGTGCATAAA | ATAGCTTTTGTTTGCCAGCATCA |
| blaCTX-M | *blaCTX-M* | Beta Lactam | CGTACCGAGCCGACGTTAA | CAACCCAGGAAGCAGGCA |
| blaTEM | *blaTEM* | Beta Lactam | CGCCGCATACACTATTCTCAG | GCTTCATTCAGCTCCGGTTC |
| blaOXA58 | *blaOXA58* | Beta Lactam | GCAATTGCCTTTTAAACCTGA | CTGCCTTTTCAACAAAACCC |
| blaKPC_3 | *blaKPC* | Beta Lactam | CAGCTCATTCAAGGGCTTTC | GGCGGCGTTATCACTGTATT |
| blaKPC_2 | *blaKPC* | Beta Lactam | GCCGCCGTGCAATACAGT | GCCGCCCAACTCCTTCA |
| blaOXA48 | *blaOXA48* | Beta Lactam | TGTTTTTGGTGGCATCGAT | GTAAMRATGCTTGGTTCGC |
| intI1_1 | *intI1* | MGE | CGAACGAGTGGCGGAGGGTG | TACCCGAGAGCTTGGCACCCA |
| oqxA | *oqxA* | MDR | GAGTCAACCTACCTCCACTATCA | GCTGCGAGTTATCCAGCAG |
| IncP_oriT | *IncP_oriT* | MGE | CAGCCTCGCAGAGCAGGAT | CAGCCGGGCAGGATAGGTGAAGT |
| IS26_1 | *IS26* | MGE | ATGGATGAAACCTACGTGAAGGTC | CGGTACTTAATCTGTCGGTGTTCA |
| ermF_1 | *ermF* | MLSB | CAGCTTTGGTTGAACATTTACGAA | AAATTCCTAAAATCACAACCGACAA |
| ermB_1 | *ermB* | MLSB | TAAAGGGCATTTAACGACGAAACT | TTTATACCTCTGTTTGTTAGGGAATTGAA |
| mphA_1 | *mphA* | MLSB | CTGACGCGCTCCGTGTT | GGTGGTGCATGGCGATCT |
| mcr1 | *mcr1* | Other | CACATCGACGGCGTATTCTG | CAACGAGCATACCGACATCG |
| qepA | *qepA* | Quinolone | GGGCATCGCGCTGTTC | GCGCATCGGTGAAGCC |
| qnrS_1 | *qnrS* | Quinolone | CCACTTTGATGTCGCAGATCTTC | CCCTCTCCATATTGGCATAGGAAA |
| sul1_2 | *sul1* | Sulfonamide | GCCGATGAGATCAGACGTATTG | CGCATAGCGCTGGGTTTC |
| sul2_2 | *sul2* | Sulfonamide | TCATCTGCCAAACTCGTCGTTA | GTCAAAGAACGCCGCAATGT |
| Enterococci | *Enterococci* | Taxonomic marker | AGAAATTCCAAACGAACTTG | CAGTGCTCTACCTCCATCATT |
| K. pneumoniae | *K. pneumoniae* | Taxonomic marker | ACGGCCGAATATGACGAATTC | AGAGTGATCTGCTCATGAA |
| Shigella | *Shigella* | Taxonomic marker | CCTTTTCCGCGTTCCTTGA | CGGAATCCGGAGGTATTGC |
| E.coli-uidA_2 | *E.coli* | Taxonomic marker | CGGAAGCAACGCGTAAACTC | TGAGCGTCGCAGAACATTACA |
| tetW | *tetW* | Tetracycline | ATGAACATTCCCACCGTTATCTTT | ATATCGGCGGAGAGCTTATCC |
| tetA_1 | *tetA* | Tetracycline | GCTGTTTGTTCTGCCGGAAA | GGTTAAGTTCCTTGAACGCAAACT |
| dfrA1_1 | *dfrA1* | Trimethoprim | GGAATGGCCCTGATATTCCA | AGTCTTGCGTCCAACCAACAG |
| dfrA8 | *dfrA8* | Trimethoprim | GGTCGCACCTGCATCGTTA | AGCGCCACCAATGACGTAG |
